# Placental derived Extracellular Matrix Supports multi-lineage cell attachment and nuclear remodeling revealed by quantitative imaging

**DOI:** 10.64898/2026.08.19.745555

**Authors:** Catalina Amurrio Zamora, Alison Ingraldi, Nilesh Dixit, Aaron J. Tabor, Fahad Mostafa

## Abstract

Decellularized extracellular matrix (dECM) scaffolds are increasingly used in regenerative medicine, yet the extent to which processed placental dECM retains properties capable of influencing cellular responses remains unclear. This study combines functional cell assays with deep learning–enabled quantitative imaging to determine how dehydrated placental ECM regulates cellular behavior across multiple human cell lineages. Human dermal fibroblasts, cardiac fibroblasts, and osteoblasts were cultured on dehydrated placental ECM or standard cell culture surfaces and assessed for cell attachment, viability, extracellular matrix production, and nuclear morphology. Placental dECM supported attachment and survival across all three cell types, while Pro-Collagen I Alpha 1 secretion varied by cell lineage relative to negative controls. To identify structural responses associated with scaffold culture, an automated imaging pipeline combining Cellpose-based nuclear segmentation with nuclear morphometric analysis was used to quantify nuclear area, eccentricity, and circularity. Quantitative profiling of hundreds of nuclei revealed scaffold-dependent remodeling of nuclear morphology that was not apparent by conventional microscopy. Cells cultured on placental dECM exhibited reduced nuclear area and increased nuclear eccentricity, while cardiac fibroblasts and osteoblasts showed alterations in nuclear circularity. These lineage-dependent morphological responses demonstrate that placental dECM provides more than a permissive substrate for cell attachment and is associated with measurable changes in cellular architecture following processing. Together, these findings support the biological relevance of processed placental dECM as a regenerative biomaterial and demonstrate the utility of quantitative single-cell morphometric analysis for detecting cell-material interactions that may not be apparent through qualitative imaging alone, guiding the rational design of regenerative therapies.

Highlights

- Placental dECM supports attachment and survival across three human cell lineages.
- Placental dECM induces lineage-dependent changes in nuclear morphology.
- Nuclear area decreases and eccentricity increases during placental dECM culture.
- Automated morphometric analysis reveals subtle cell-biomaterial responses.

**Graphical abstract:** 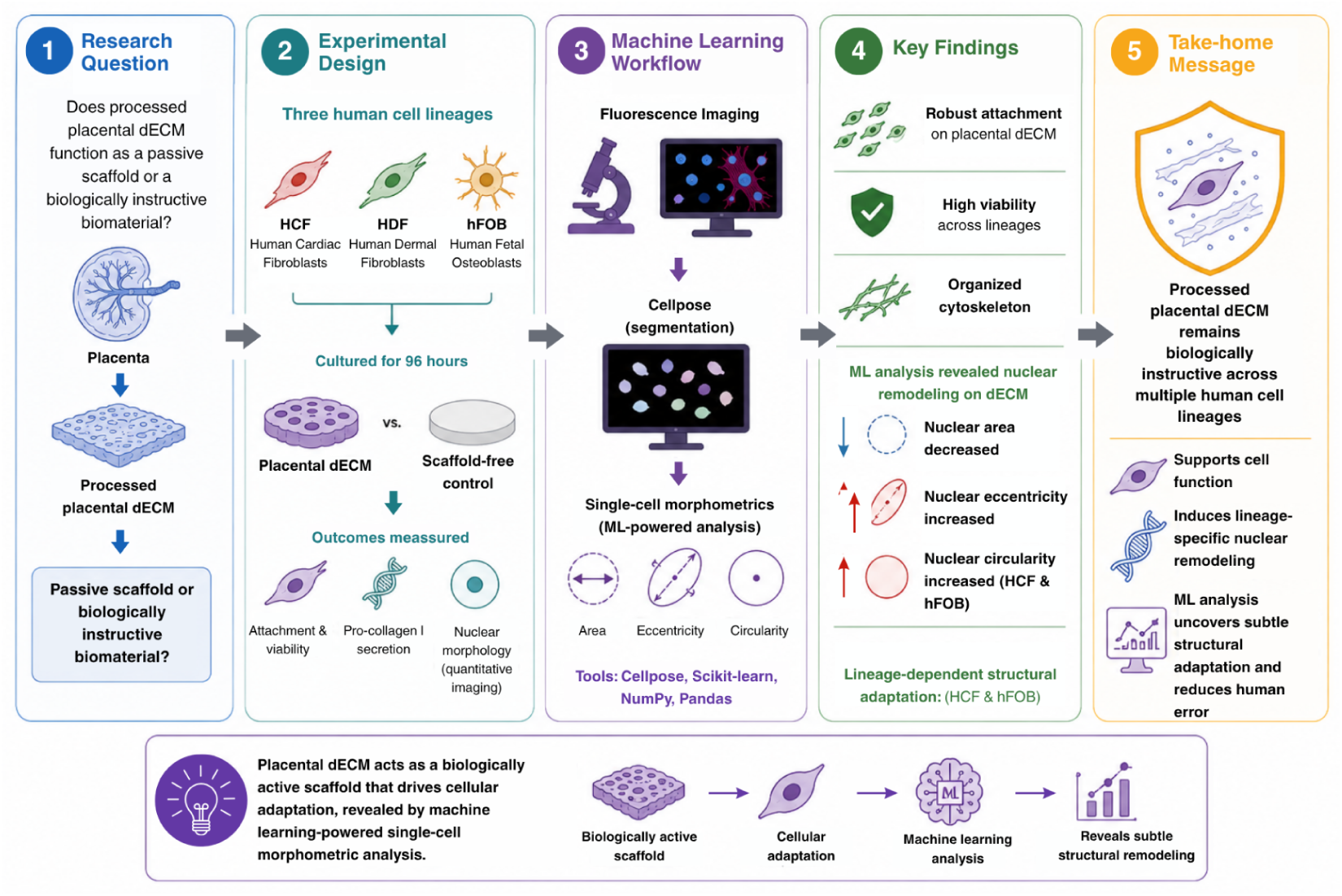

## Introduction

The extracellular matrix (ECM) is a dynamic network of proteins, glycoproteins, proteoglycans, and matrix-bound signaling molecules that regulate fundamental aspects of cellular behavior, including adhesion, survival, proliferation, migration, and differentiation [1,2,3,4]. Besides supplying mechanical support, the ECM functions as an active regulator of tissue homeostasis and repair by modulating cell-matrix interactions as well as controlling the availability of growth factors within the cellular microenvironment [5,6,7,8]. Given these characteristics, ECM-derived biomaterials have become an important component of regenerative medicine strategies aimed at promoting tissue repair and functional recovery.

Decellularized extracellular matrix (dECM) scaffolds have shown clinical utility across a wide range of applications, including wound healing, soft tissue reconstruction, and cardiovascular repair [9]. These biomaterials can preserve structural and biochemical features of native tissues, which provide a microenvironment that is capable of supporting cellular attachment, viability, and remodeling. Nevertheless, ECM composition is inherently tissue specific, which reflects the unique structural and signaling requirements of different organs and cell populations [1]. For instance, bone ECM contains a specialized matrix that is rich in collagen and mineral-associated proteins that can regulate osteogenic differentiation and tissue mineralization [10,11], while cardiac ECM is rich in structural proteins that support myocardial architecture and contribute to tissue repair following injury [12,13]. These observations suggest that ECM composition plays a critical role in directing lineage-specific cellular responses.

The biological activity of ECM scaffolds is influenced not only by methods used during processing and preservation but also by the tissue from which the matrix is derived. Native ECM composition varies substantially among tissues, and these biochemical and structural differences contribute to tissue-specific regulation of cellular behavior [1]. Clinical ECM products commonly undergo decellularization and dehydration procedures to improve product shelf-life stability and facilitate handling. Although these approaches are intended to preserve matrix architecture, they may alter the activity of the signaling molecules associated with the matrix that contribute to cell-guiding behavior [14]. Therefore, an important unanswered question is whether processed ECM scaffolds retain lineage-specific biological cues or primarily serve as permissive substrates that support cell attachment and survival, regardless of cell type.

This question is particularly relevant for placental-derived ECM biomaterials. In recent years, placental tissues have emerged as potential scaffold sources mainly due to their availability, biocompatibility, immunomodulatory properties, and rich extracellular matrix composition [15,16]. Placental ECM biomaterials are increasingly used across a wide range of regenerative medicine applications, often applied across tissues with distinct cellular and ECM requirements. However, relatively little is known about how a single, processed and dehydrated placental ECM scaffold interacts differentially with distinct human cell lineages, or the extent to which tissue-instructive properties are preserved following processing.

Recent advances in computational image analysis provide an opportunity to characterize cellular responses to biomaterials quantitatively. Automated image segmentation and high-content imaging approaches enable objective extraction of morphological features from fluorescence microscopy datasets, which allows the detection of subtle phenotypic differences at the single-cell level [17,18,19]. When combined with biochemical measurements, these methods can provide a quantitative framework for evaluating cell-material interactions and assessing cellular adaptation to engineered microenvironments [20,21].

This study investigated whether a dehydrated placenta-derived dECM functions as a tissue-instructive scaffold or as a broadly permissive substrate across multiple human cell lineages. Human dermal fibroblasts (HDFa), Cardiac Fibroblasts (HCF), and Osteoblasts (hFOB) were evaluated using viability assays, collagen production, and deep learning single-cell morphometric analysis. Placental dECM supported attachment and viability across all cell types while quantitative imaging identified scaffold-dependent nuclear remodeling that was not apparent by conventional microscopy. These findings indicate that processed placental dECM retains biologically instructive properties while demonstrating the value of high-content morphometric profiling for biomaterial evaluation.

## Materials and Methods

### ECM Scaffold Preparation

Cell culture and scaffold preparation procedures were adopted from previously reported methods [22] and are described in full below to ensure reproducibility. The ECM scaffolds used in this study were donated by an FDA-registered tissue bank that offers commercially available dehydrated placental membranes, Axolotl Graft™ 2cm x 4cm (Axolotl Biologix, Scottsdale, AZ, USA) (Fig. 1). The material was previously processed, decellularized, and terminally irradiated while preserving the native ECM architecture and composition [14].

**Figure 1:**
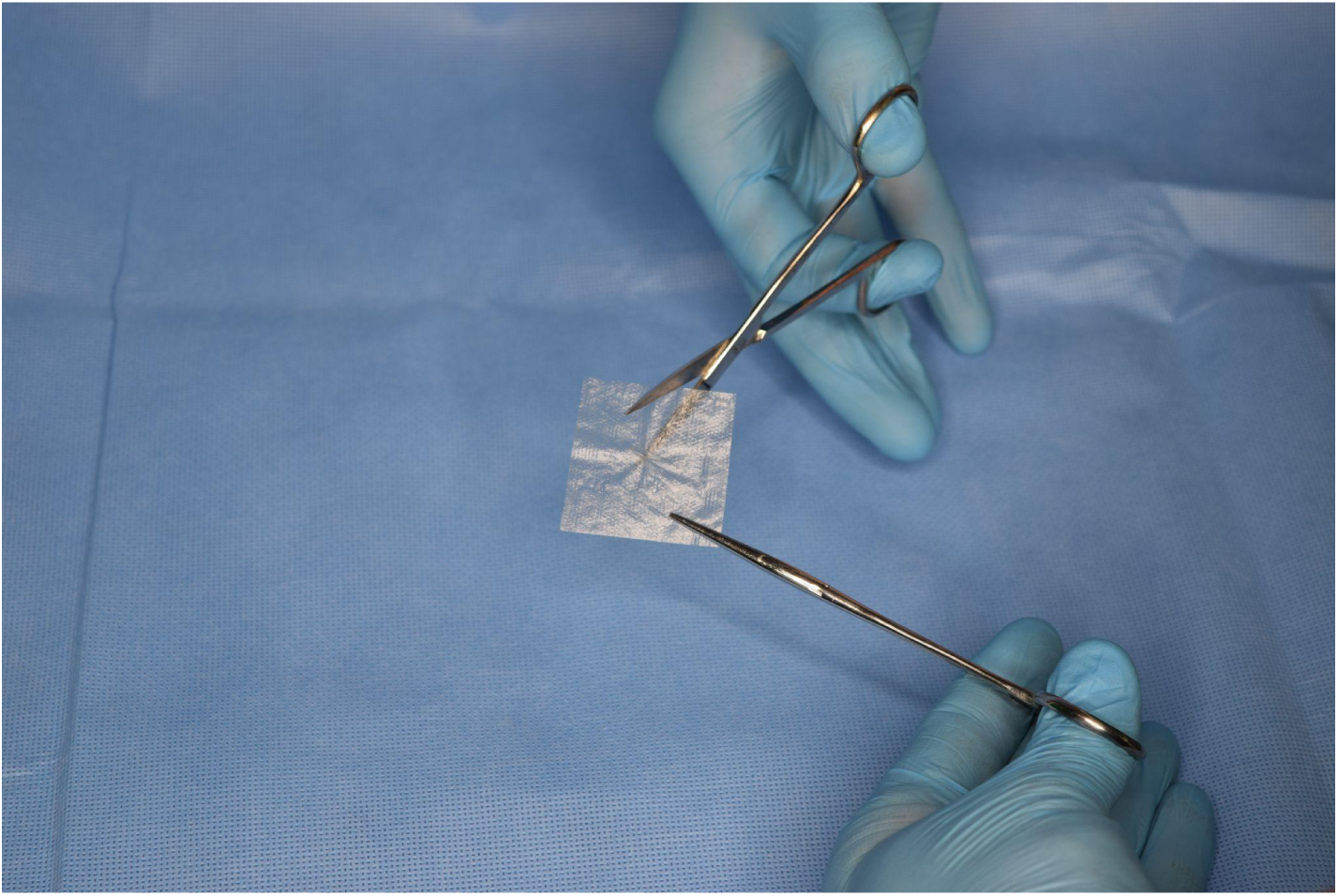
Representative image of a dehydrated placenta-derived ECM scaffold.

Under aseptic conditions, membranes were sectioned into circular biopsies using a 10 mm biopsy punch (Robbins Instruments, Houston, TX, USA) and placed into 24-well plates (Costar Multiple Well Plate, 24-well with Lid. Corning CellBIND surface Polystyrene; Ref 3337; Corning, NY, USA). Before cell seeding, membranes were hydrated for 24 hours in the respective culture media for each cell type (HCF, HDF, and hFOB). Wells containing hydrated ECM scaffolds served as the experimental scaffold conditions, while wells without membranes served as tissue culture controls, and membrane-only wells containing no cells served as the negative controls.

### Cell Culture

Human Dermal Fibroblasts (HDFa; PCS-201-012, ATCC, Manassas, VA, USA), Human Cardiac Fibroblasts (HCF; adult ventricular, 306V-05a, Cell Applications Inc., San Diego, CA, USA), and Human Fetal Osteoblasts (hFOB 1.19; CRL-11372, ATCC, Manassas, VA, USA) were cultured according to manufacturer protocols. HDF cells were maintained in Dulbecco’s Modified Eagle Medium (DMEM; ATCC®, Manassas, VA, USA; Cat. No. 30-2002) supplemented with 1% Penicillin-Streptomycin (Sigma-Aldrich, St. Louis, MO, USA; Cat. No. P4083), HCF cells were maintained in manufacturer-provided growth medium (Cell Applications Inc., San Diego, CA, USA), and hFOB 1.19 cells were maintained in a 1:1 mixture of Ham’s F-12 Medium and DMEM supplemented according to manufacturer recommendations (Sigma-Aldrich, St. Louis, MO, USA; Cat. No. C27001). HDF and hFOB cells were expanded from passage 2 to passage 3, whereas HCF cells were expanded from passage 1 to passage 2 before experimental seeding.

HCF and hFOB cells were cultured in T75 CellBIND flasks (Corning Inc., Corning, NY, USA), while HDF cells were cultured on T75 Cell Culture Flask Polystyrene nonpyrogenic (Corning Inc., Corning, NY, USA) and maintained in a VWR Air Jacketed CO_2_ Incubator (Avantor, Radnor, PA, USA) at 37°C and 5% CO_2_ until approximately 80% confluence. Culture medium was replaced 24 hours after initial seeding and subsequently at 72 hours post-seeding. Cells were detached using TrypLE™ Select (Gibco, Thermo Fisher Scientific, Waltham, MA, USA), neutralized with growth medium, centrifuged at 1200rpm for 5 minutes, and resuspended in fresh culture medium. Cell concentration and viability were determined by 0.4% Trypan Blue (Gibco, Thermo Fisher Scientific, Waltham, MA, USA; Cat. No. T10282) using a Countless™ 3 Automated Cell Counter (Thermo Fisher Scientific, Waltham, MA, USA).

### Experimental Seeding

Cell suspensions were prepared for experimental seeding using manufacturer-recommended seeding densities for the placental ECM scaffold. HDF, HCF, and hFOB 1.19 cells were seeded at densities of 3,000 cells/cm^2^, 7,000 cells/cm^2^, and 5,000 cells/cm^2^, respectively. Experimental groups consisted of cells seeded onto hydrated placenta-derived ECM scaffolds, and positive control groups consisted of cells seeded directly into wells without an ECM scaffold. Negative controls contained only hydrated ECM scaffolds. Each experimental condition was performed in triplicate across two independent 24-well plates.

For ECM scaffold conditions, cells were seeded onto hydrated ECM scaffolds as described above. A 2.5 mL volume of cell suspension was gently pipetted onto the center of each scaffold to promote uniform cell distribution. Care was taken to avoid disturbing the scaffold and to prevent cells from being introduced beneath it. Plates were incubated under standard culture conditions (37°C, 5% CO₂). At 24 hours post-seeding, 0.5 mL of the appropriate growth medium was added to each well. After 48 hours, the culture medium was gently aspirated and replaced with fresh medium to maintain optimal growth conditions and promote stable cell attachment to the ECM scaffolds. Cultures were maintained for a total of 96 hours before downstream analyses, including fluorescence imaging, quantitative morphometric analysis, cell viability assessment, and Pro-Collagen I Alpha 1 quantification.

### Pro-Collagen I Alpha 1 Quantification

Extracellular Pro-Collagen I Alpha 1 production was quantified using a Human Pro-Collagen I Alpha 1 SimpleStep ELISA kit (Abcam, Cambridge, UK; Cat. No. ab210966). Conditioned media samples from HDF, HCF, and hFOB cultures were collected following 96 hours of incubation were centrifuged at 2,000 rpm for 10 minutes and diluted 1:4 in sample diluent prior to analysis.

A standard curve ranging from 0 to 2000 pg/mL was prepared using serial dilutions of the supplied protein standard. Standards and diluted samples were analyzed in technical triplicate in a 96-well microplate. Signal development was monitored kinetically at 600 nm for 10 min using a Synergy™ H1 Hybrid Multi-Mode Microplate Reader (BioTek Instruments, Winooski, VT, USA). The reaction was terminated with stop solution, and absorbance was measured at 450 nm. Raw absorbance values were exported to Microsoft Excel (Microsoft Corp., Redmond, WA, USA), where a standard curve was generated and Pro-Collagen I Alpha 1 concentrations were determined by interpolation.

### Fluorescence Staining and Imaging

At the experimental endpoint (96 hours post-seeding), media was removed, and samples were gently rinsed twice with HBSS and fixed with 4% paraformaldehyde (Thermo Fisher Scientific, Waltham, MA, USA; Cat. No. J61984.AP) for 15 minutes at room temperature, The paraformaldehyde solution was removed, and samples were washed twice with HBSS and stored in HBSS at 4°C until staining.

All cells were labeled to visualize the nuclei and evaluate cellular morphology and cytoskeletal organization. Cell nuclei were first stained with 300 nM 4’-6-diamidino-2-phenylindole (DAPI; Thermo Fisher Scientific, Waltham, MA, USA; Cat. No. 62248) for 15 minutes, followed by washing with Hank’s Balanced Salt Solution (HBSS; Thermo Fisher Scientific, Waltham, MA, USA; Cat. No. 14175095). Filamentous actin (F-actin), a major structural component of the actin cytoskeleton, was subsequently labeled to visualize cell shape and cytoskeletal organization using Alexa Fluor 488-conjugated phalloidin (Thermo Fisher Scientific, Waltham, MA, USA; Cat. No. A12379). Samples were incubated with phalloidin for 30 minutes at room temperature in the dark then washed with HBSS.

Fluorescence images were acquired using a Leica DMi8 fluorescence microscope (Leica Microsystems, Wetzlar, Germany) equipped with appropriate filters for DAPI and Alexa Fluor 488 detection. Images were collected using a 20x objective lens, corresponding to a total magnification of 200x. Multiple representative fields of view were acquired from each sample while avoiding scaffold edges and visible artifacts. Exposure time and illumination settings were held constant across all experimental conditions to ensure comparability between groups.

### Automated Image Analysis

Fluorescence microscopy images were processed using a computational image analysis pipeline developed in Python (version 3.13.5). To improve computational efficiency and segmentation performance, each full-field image was divided into 12 equally sized, non-overlapping tiles without altering the original fluorescence and intensity data. Approximately 12,000 image tiles were generated from the complete imaging dataset and processed individually.

Nuclear segmentation was performed using Cellpose (version 3.1), a deep learning-based framework designed for robust biological image segmentation [18,23] For each image tile, the DAPI fluorescence channel was extracted and used as input to the pretrained nuclei model. This model generated labeled segmentation masks that detected objects corresponding to a single nucleus, which were subsequently used to quantify nuclear counts and extract morphometric measurements for downstream analysis.

### Segmentation Validation and Morphometric Feature Extraction

To evaluate segmentation performance, a validation dataset that consisted of 900 image tiles was randomly assembled by selecting 100 tiles from each of the nine experimental groups. The automated nuclear segmentations generated by Cellpose were compared with manually annotated ground-truth masks created for each validation image. Segmentation accuracy was quantified using the Dice Similarity Coefficient (DSC) and Intersection-over-Union (IoU), both of which measure the spatial overlap between automated segmentations and manually annotated reference masks. In order to assess segmentation quality across a range of performance levels, two low-scoring, two intermediate-scoring, and two high-scoring image tiles, based on their DSC and IoU values, were selected from each experimental group for detailed manual review, resulting in a final validation cohort of 54 image tiles.

Quantitative morphometric features were extracted from the segmentation masks using the regionprops_table function implemented in the scikit-image library [24]. Measurements were calculated on a per-nucleus basis and included nuclear area, perimeter, and eccentricity, where the last one refers to the measure of nuclear elongation ranging from 0 for a perfect circle to values approaching 1 for increasingly elongated ellipses. Circularity was calculated according to the equation:

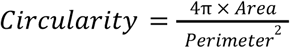

where Area represents the segmented nuclear area, and Perimeter represents the perimeter of the corresponding segmented nucleus. Circularity values approaching 1 indicate increasingly circular nuclei. All morphometric measurements were compiled into a single-cell dataset in which each observation represented an individual nucleus and was annotated according to cell type and experimental condition. These quantitative features were subsequently used to compare nuclear morphology between the ECM scaffold and control groups.

## Results

### Placenta-derived ECM scaffolds support attachment of multiple human cell lineages

DAPI and phalloidin staining demonstrated successful attachment and survival across all three cell types cultured on placental ECM scaffolds and tissue culture plastic controls (Fig. 2).

**Figure 2.**
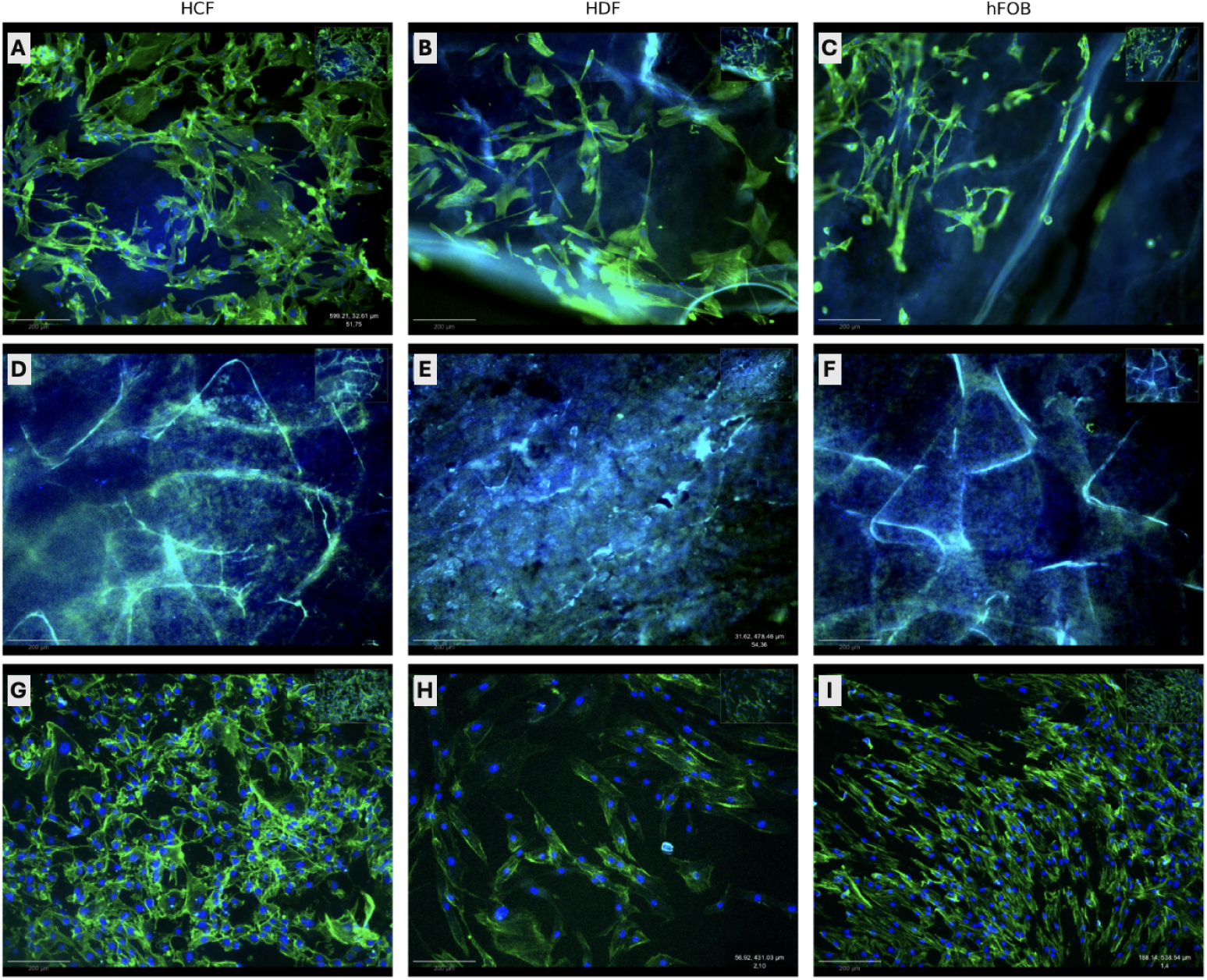
Representative fluorescence microscopy of human cell lineages cultured on dehydrated placental dECM scaffolds. Human Cardiac Fibroblasts (HCF), Human Dermal Fibroblasts (HDF), and Human Fetal Osteoblasts (hFOB) were stained with DAPI (blue) to visualize nuclei and phalloidin (green) to visualize F-actin. Panels A-C show each lineage adherent to the placental dECM scaffold, demonstrating attachment and cytoskeletal organization along the scaffold surface. Panels D-F show acellular scaffold controls incubated in the corresponding culture media, confirming the absence of cells while illustrating the architecture of the scaffold. Panels G-I show cells cultured without scaffolds, serving as positive controls for normal cellular morphology and organization. Reproduced from Amurrio Zamora [22].

Cells that were cultured on tissue culture plastic plate wells formed relatively uniform monolayers with well-defined actin cytoskeletal organization and evenly distributed nuclei (Fig 2G - I). Correspondingly, cells seeded onto placental ECM scaffolds exhibited extensive cytoskeletal structures and clear nuclear staining, which indicated that the scaffold provided a suitable substrate for cellular attachment and early spreading (Fig 2A - C). While differences in cellular organization and distribution were visually apparent between ECM and control conditions, qualitative microscopy alone was insufficient to objectively characterize these changes.

Negative control wells that contained acellular ECM scaffolds did not show detectable nuclear or cytoskeletal staining, confirming that fluorescent signals observed in experimental samples originated from attached cells rather than background scaffold fluorescence (Fig 2D - F).

### Automated image segmentation enables quantitative morphometric analysis

To quantify the cellular responses to the scaffold, fluorescence microscopy images were processed using an automated image-analysis workflow based on the Cellpose deep learning segmentation framework. DAPI fluorescence images were partitioned into equal-sized, smaller tiles before segmentation, which generated approximately 12,000 image tiles for downstream analysis.

Segmentation accuracy was evaluated by comparing masks generated by Cellpose with the manually annotated reference masks using the DSC and IoU. Positive control samples consistently demonstrated the highest agreement between automated and manual annotations with mean DSC scores ranging from 0.636 to 0.833 and mean IoU values ranging from 0.584 to 0.783 across all three cell types (Table 1). Additionally, hFOB positive controls achieved the highest segmentation accuracy, with mean DSC and IoU values of 0.833 and 0.783, respectively. ECM samples showed intermediate agreement (DSC: 0.380 - 0.486; IoU: 0.330 - 0.399), which reflected the increased complexity of segmenting nuclei within the scaffold microenvironment. Negative controls produced DSC and IoU values of zero or near zero. This indicated the absence of meaningful segmentation. The non-zero values observed specifically for the HCF negative control (DSC: 0.074; IoU: 0.058) likely reflect occasional background segmentation rather than true nuclear detection.

**Table 1.**
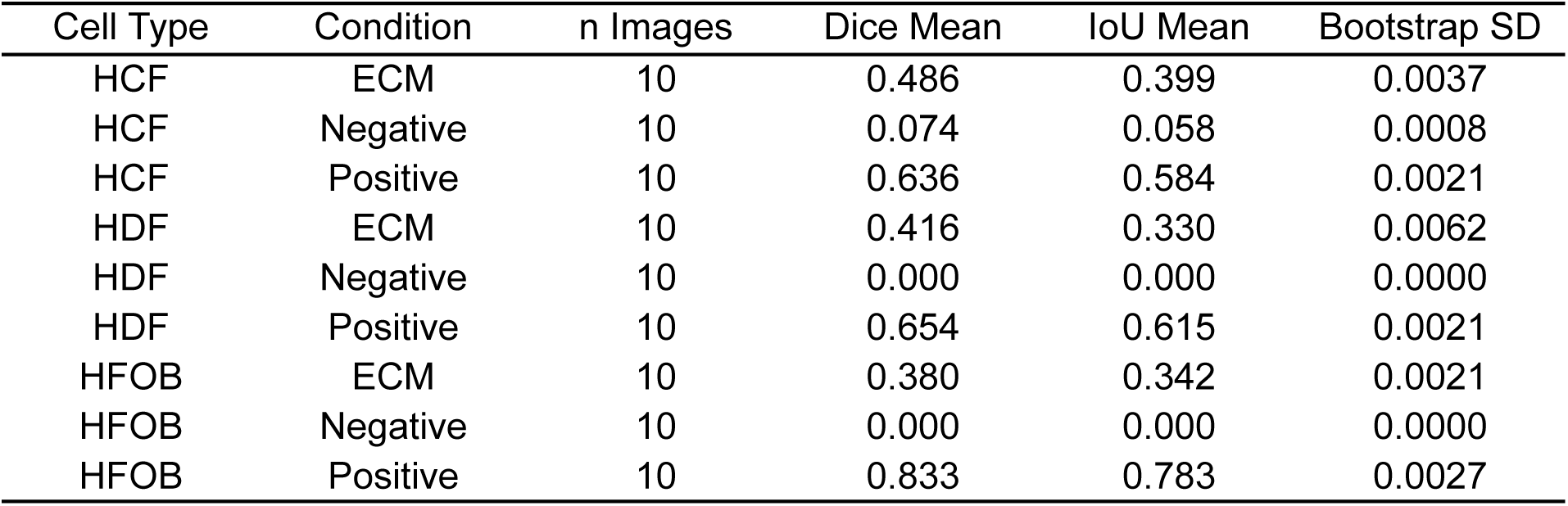
Segmentation Performance Across Cell Types and Culture Conditions. Performance of the automated Cellpose-based image segmentation pipeline across three human cell types (HCF, HDF, and HFOB) cultured under extracellular matrix (ECM), positive-control, and negative-control conditions. Segmentation accuracy was evaluated using 10 microscopy images per condition. Mean DSC and IoU quantify the agreement between predicted segmentation masks and manually annotated ground-truth masks, while the bootstrap standard deviation (Bootstrap SD) measures the robustness and reproducibility of the segmentation results. Positive-control images consistently yielded the highest segmentation accuracy, whereas negative controls showed negligible overlap with the ground truth. ECM samples demonstrated intermediate but reproducible performance across all cell types, supporting the reliability of the proposed segmentation pipeline for quantitative morphometric analysis. Adapted from Amurrio Zamora [22].

| Cell Type | Condition | n Images | Dice Mean | IoU Mean | Bootstrap SD |
| --- | --- | --- | --- | --- | --- |
| HCF | ECM | 10 | 0.486 | 0.399 | 0.0037 |
| HCF | Negative | 10 | 0.074 | 0.058 | 0.0008 |
| HCF | Positive | 10 | 0.636 | 0.584 | 0.0021 |
| HDF | ECM | 10 | 0.416 | 0.330 | 0.0062 |
| HDF | Negative | 10 | 0.000 | 0.000 | 0.0000 |
| HDF | Positive | 10 | 0.654 | 0.615 | 0.0021 |
| HFOB | ECM | 10 | 0.380 | 0.342 | 0.0021 |
| HFOB | Negative | 10 | 0.000 | 0.000 | 0.0000 |
| HFOB | Positive | 10 | 0.833 | 0.783 | 0.0027 |

In contrast, cells cultured on ECM scaffolds showed greater variability in segmentation performance, which was reflected by broader distributions of DSC and IoU values (Fig. 3; Fig. 4). The increased variability observed was consistent with a greater heterogeneity in nuclear morphology and spatial organization observed in scaffold-grown cells. Negative controls produced near-zero segmentation scores because no nuclei were present in those samples. Representative examples of manually annotated ground-truth masks, Cellpose-generated segmentations, and overlay images are shown in Fig. 5. The close agreement between manual and automated segmentations across all experimental conditions supports the accuracy and robustness of the segmentation pipeline for subsequent quantitative morphometric analysis.

**Figure 3.**
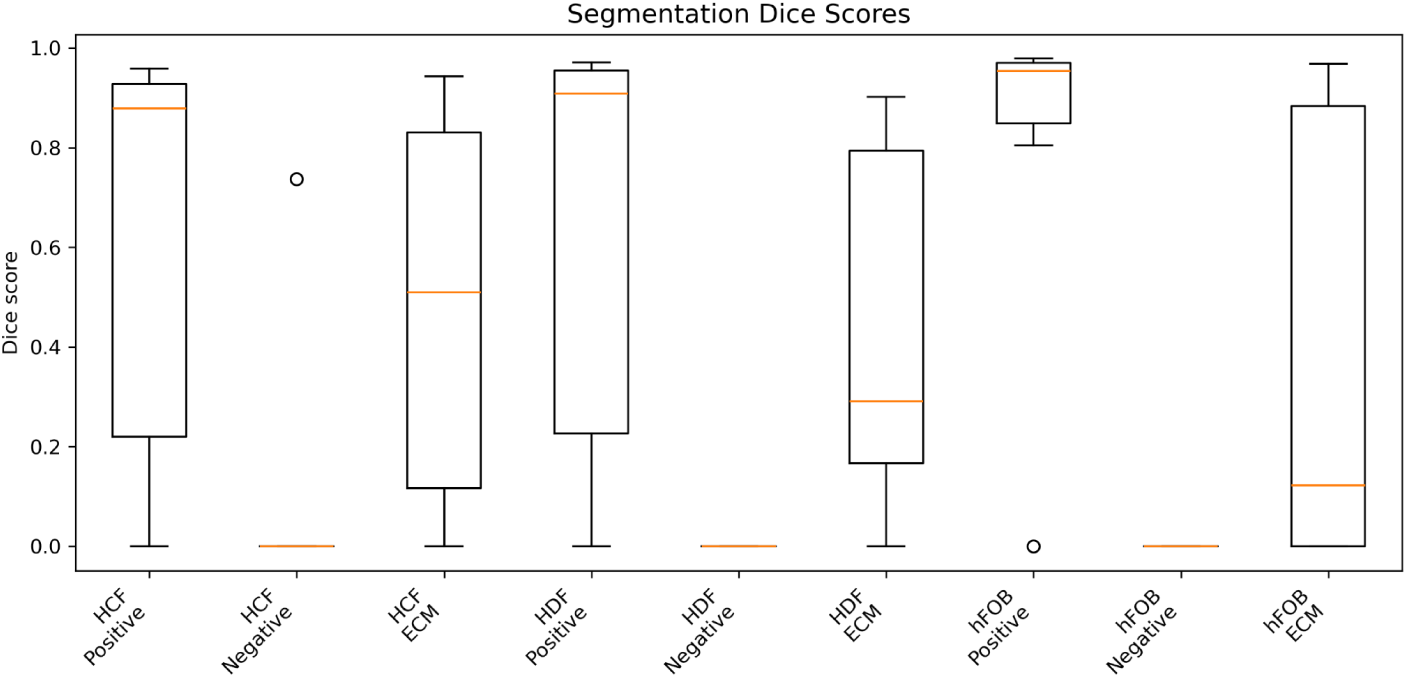
Distribution of Dice similarity coefficients (DSC) across cell types and experimental conditions. Boxplots summarize the distribution of DSC similarity coefficients obtained from automated Cellpose-based segmentation for HCF, HDF, and hFOB cultured under positive-control, ECM, and negative-control conditions. Boxes represent the interquartile range (IQR), the orange line indicates the median, whiskers extend to 1.5 × IQR, and circles denote outliers. Reproduced from Amurrio Zamora [22].

**Figure 4.**
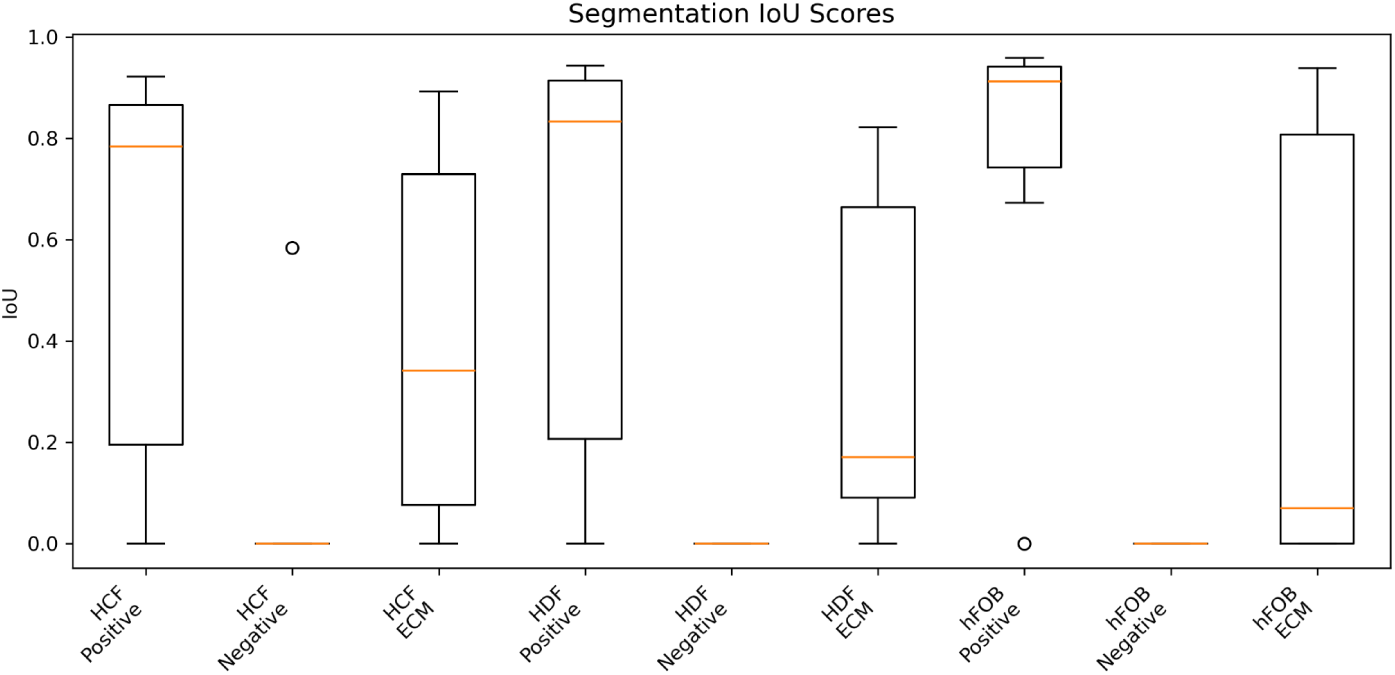
Distribution of Intersection over Union (IoU) scores across cell types and experimental conditions. Boxplots show the distribution of IoU values for automated Cellpose-based segmentation across HCF, HDF, and HFOB cultures under positive-control, ECM, and negative-control conditions. Higher IoU values indicate greater agreement between predicted segmentation masks and manually annotated ground-truth masks. Boxes represent the interquartile range (IQR), the center line indicates the median, whiskers extend to 1.5 × IQR, and circles denote outliers. Reproduced from Amurrio Zamora [22].

**Figure 5.**
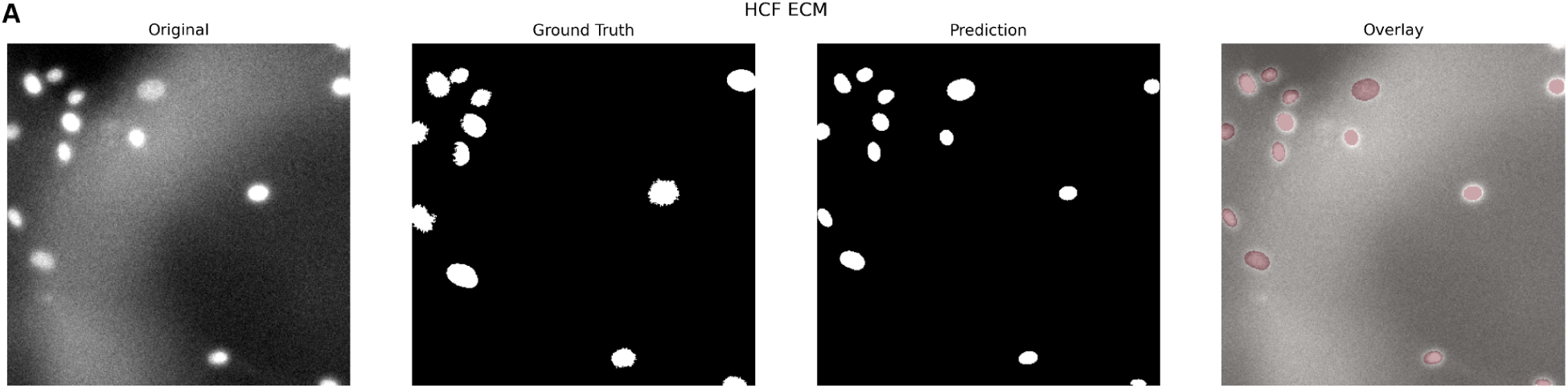
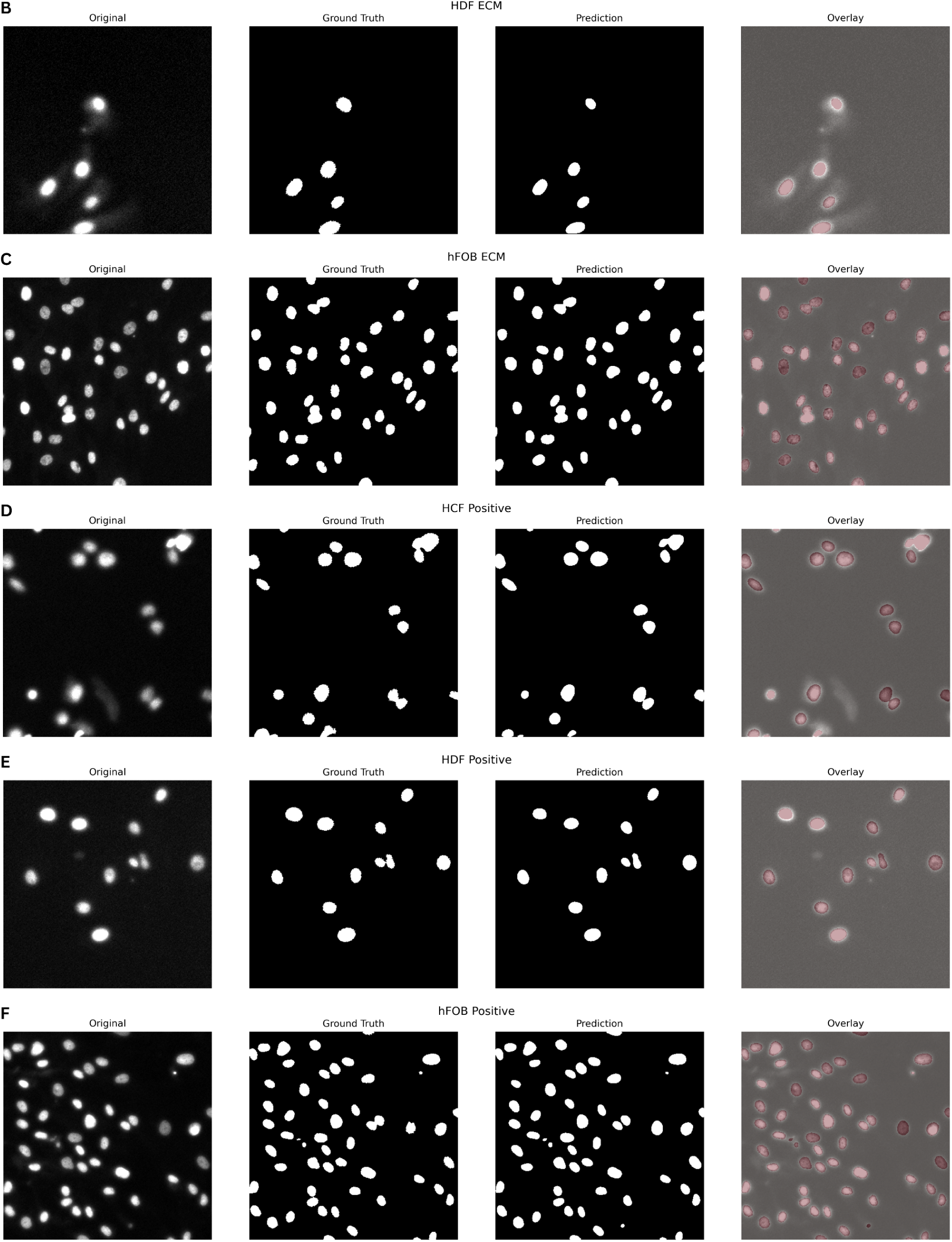
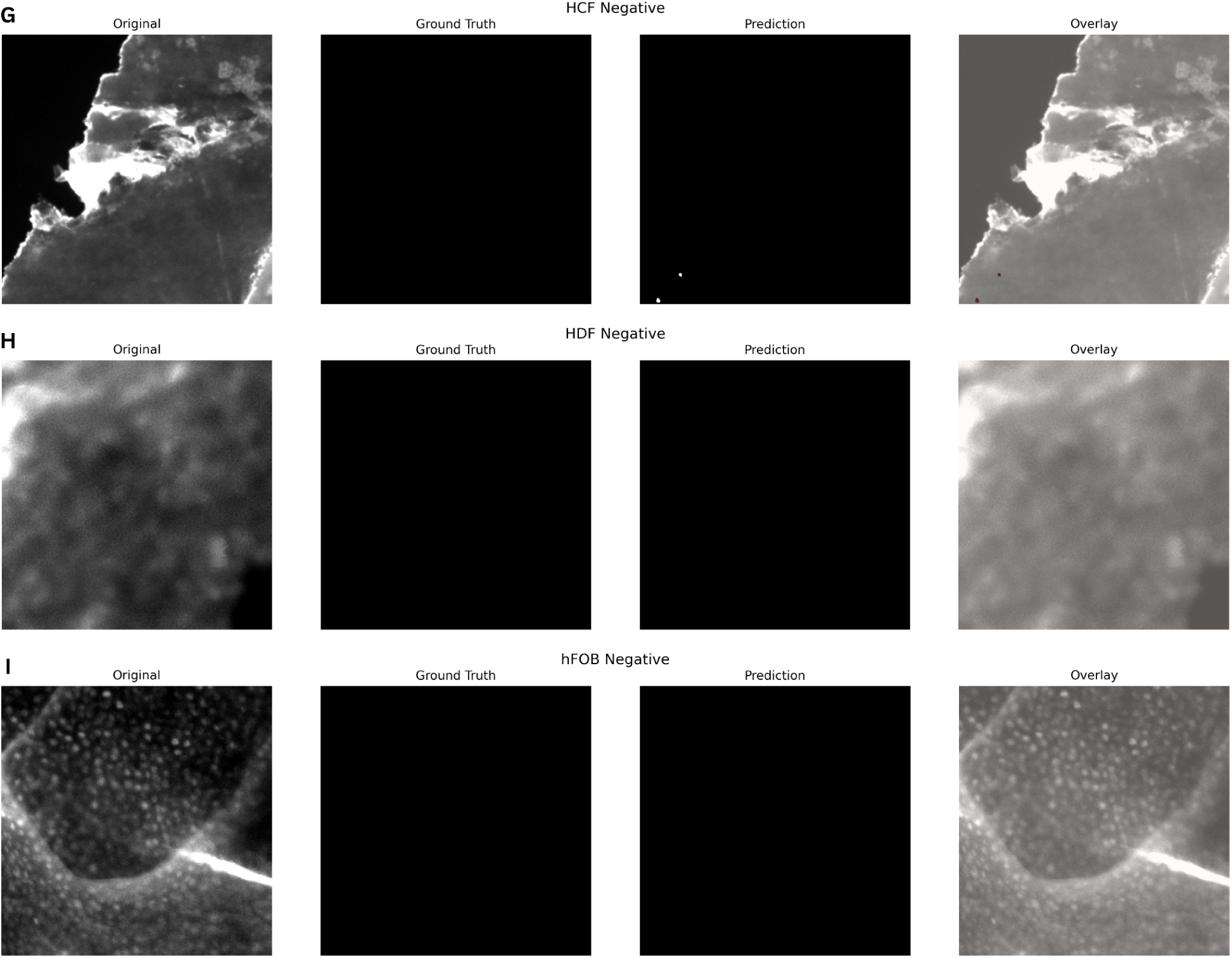
Representative examples of automated nuclear segmentation across experimental conditions. Representative DAPI fluorescence images from HCF, HDF, and hFOB cultured on placental dECM (A-C), cells cultured without scaffold (D-F), and acellular scaffold controls (G-I). For each condition, the original fluorescence image, manually annotated ground truth mask, Cellpose prediction, and overlay of predicted masks on the original image are shown. The close agreement between manual annotations and automated predictions demonstrates accurate nuclear segmentation across diverse cellular morphologies, while negative controls show minimal false-positive detections. Reproduced from Amurrio Zamora [22].

### Placental ECM alters nuclear morphology across multiple human cell lineages

Following segmentation, individual nuclei were analyzed to quantify morphological responses to the ECM scaffold. A total of 509 nuclei were identified and measured across all three cell line cultures under ECM and control conditions. For each nucleus, quantitative features describing nuclear size and shape, including area, circularity, and eccentricity, were extracted.

Placental dECM significantly reduced nuclear area across all three cell types when compared to positive controls (Fig. 6A). Overall, cells cultured on standard tissue culture plastic exhibited larger nuclear areas than cells cultured on ECM scaffolds, indicating altered nuclear morphology relative to positive controls.

**Figure 6.**
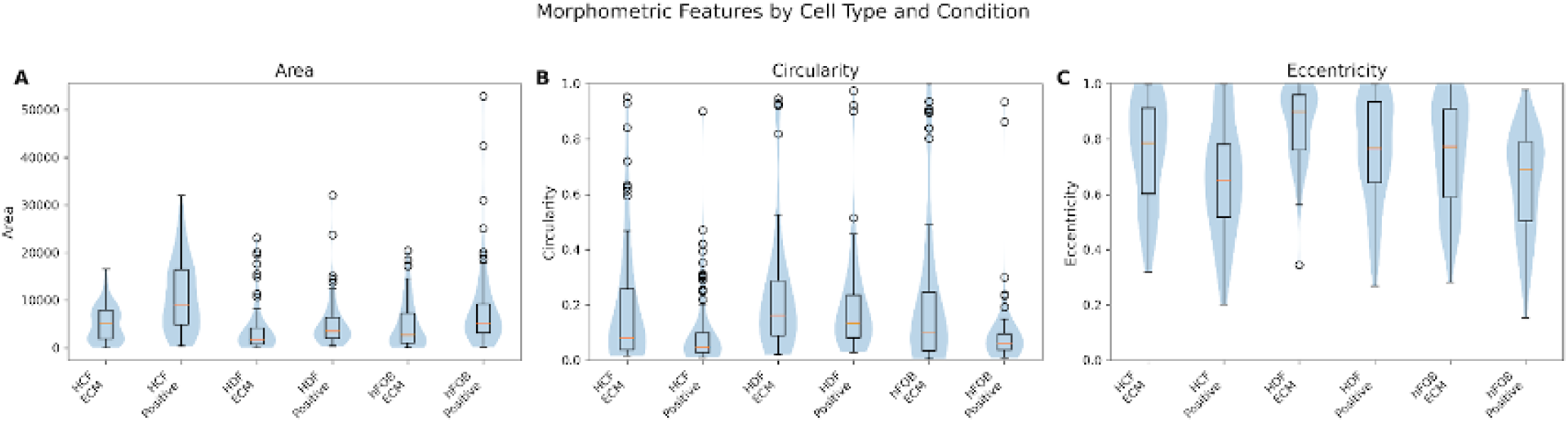
Quantitative morphometric analysis identifies scaffold-dependent changes in nuclear morphology across multiple human cell lineages. Distribution of nuclear area (A), circularity (B), and eccentricity (C) measured following automated Cellpose segmentation and single-cell morphometric analysis. HCF, HDF, and hFOB cultured on placental dECM were compared with cells cultured without an ECM scaffold. Violin plots show the distribution of the individual nuclei, while embedded boxplots indicate the median and interquartile range. Placental dECM was associated with reduced nuclear area and increased eccentricity across all three cell lineages, with significant alterations in circularity observed in HCF and hFOB. Reproduced from Amurrio Zamora [22].

Nuclear shape was altered by placental dECM, producing increased eccentricity across all three lineages and significant changes in circularity in HCF and hFOB cells (Fig, 6B - C). Cells cultured on placental ECM scaffolds showed increased circularity and eccentricity relative to their corresponding controls; this indicated alterations in nuclear architecture associated with scaffold interaction.

For HCF cells, median circularity decreased from 0.045 under control conditions to 0.0079 on ECM scaffolds. Likewise, hFOB cells demonstrated an increase from 0.062 to 0.103, whereas HDF cells showed a more modest increase from 0.136 to 0.162. Nuclear eccentricity was consistently larger across all ECM conditions, increasing from 0.652 to 0.785 for HCF, from 0.766 to 0.898 for HDF, and from 0.692 to 0.773 for hFOB cells.

Statistical comparisons were performed using the Mann-Whitney U test. Circularity differences reached statistical significance for HCF (p = 2.02 × 10⁻⁴) and hFOB (p = 4.58 × 10⁻³), whereas differences in HDF cells were not significant (p = 0.374). In contrast, eccentricity differed significantly between ECM and control conditions across all three cell types, with p-values of 4.83 × 10⁻³, 1.80 × 10⁻³, and 5.82 × 10⁻⁴ for HCF, HDF, and hFOB cells, respectively (Table 2).

**Table 2.**
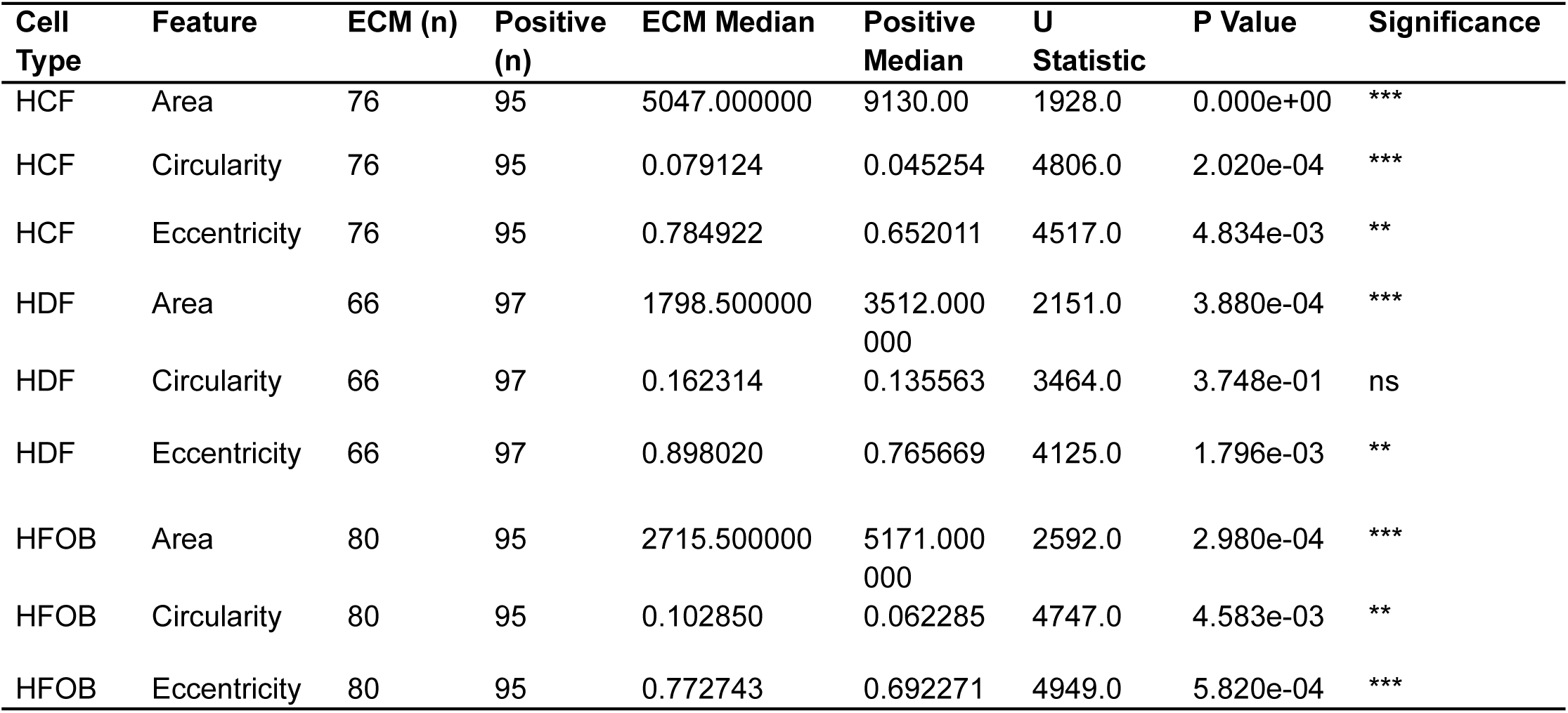
Statistical comparison of nuclear morphometric features between placental dECM and positive-control cultures. Mann–Whitney U tests were performed to compare nuclear area, circularity, and eccentricity between cells cultured on placenta dECM and positive control for HCF, HDF, and hFOB. The table reports sample size (*n*), median feature values, Mann–Whitney U statistics, *P* values, and significance levels. Significant differences in nuclear area and eccentricity were observed across all three cell types. In contrast, circularity differed significantly in HCF and HFOB but not in HDF, indicating cell-type-dependent nuclear remodeling induced by the dECM microenvironment. Adapted from Amurrio Zamora [22].

| Cell Type | Feature | ECM (n) | Positive (n) | ECM Median | Positive Median | U Statistic | P Value | Significance |
| --- | --- | --- | --- | --- | --- | --- | --- | --- |
| HCF | Area | 76 | 95 | 5047.000000 | 9130.00 | 1928.0 | 0.000e+00 | *** |
| HCF | Circularity | 76 | 95 | 0.079124 | 0.045254 | 4806.0 | 2.020e-04 | *** |
| HCF | Eccentricity | 76 | 95 | 0.784922 | 0.652011 | 4517.0 | 4.834e-03 | ** |
| HDF | Area | 66 | 97 | 1798.500000 | 3512.000<br>000 | 2151.0 | 3.880e-04 | *** |
| HDF | Circularity | 66 | 97 | 0.162314 | 0.135563 | 3464.0 | 3.748e-01 | ns |
| HDF | Eccentricity | 66 | 97 | 0.898020 | 0.765669 | 4125.0 | 1.796e-03 | ** |
| HFOB | Area | 80 | 95 | 2715.500000 | 5171.000<br>000 | 2592.0 | 2.980e-04 | *** |
| HFOB | Circularity | 80 | 95 | 0.102850 | 0.062285 | 4747.0 | 4.583e-03 | ** |
| HFOB | Eccentricity | 80 | 95 | 0.772743 | 0.692271 | 4949.0 | 5.820e-04 | *** |

Collectively, these findings indicate that although the dehydrated placental ECM scaffold supported attachment across multiple human cell lineages, it also induced measurable changes in nuclear morphology that varied according to cell type.

To determine whether cellular interaction with the scaffold was accompanied by ECM protein production, conditioned media were analyzed using a kinetic Pro-Collagen I Alpha 1 ELISA assay. A standard curve was generated using known protein concentrations and signal development rate, confirming assay suitability for quantitative analysis (Fig. 7).

**Figure 7.**
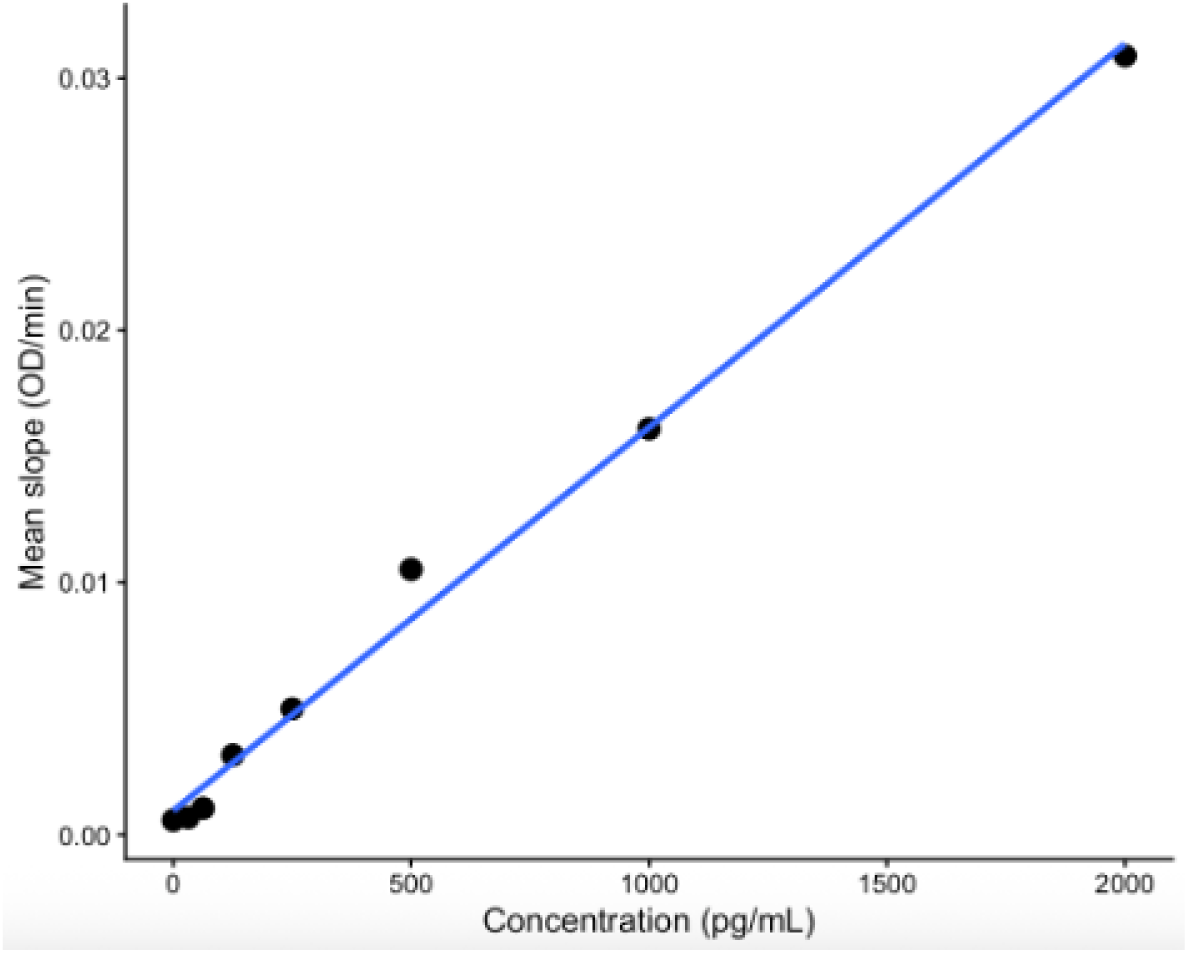
Standard curve for kinetic ELISA quantification of Pro-Collagen I Alpha 1. Mean reaction slopes (OD/min) were plotted against known Pro-Collagen I Alpha 1 concentrations to generate the calibration curve used for sample quantification. The blue line represents the linear regression fit. Reproduced from Amurrio Zamora [22].

Detectable Pro-Collagen I Alpha 1 expression was observed across all three cell types under both ECM and control conditions. Positive control cultures exhibited higher mean kinetic slopes than scaffold-grown cultures, with values of 0.330 ± 0.005 OD/min versus 0.207 ± 0.021 OD/min for hFOB cells, 0.123 ± 0.014 OD/min versus 0.117 ± 0.054 OD/min for HCF cells, and 0.192 ± 0.028 OD/min versus 0.166 ± 0.012 OD/min for HDF cells (Fig. 8).

**Figure 8.**
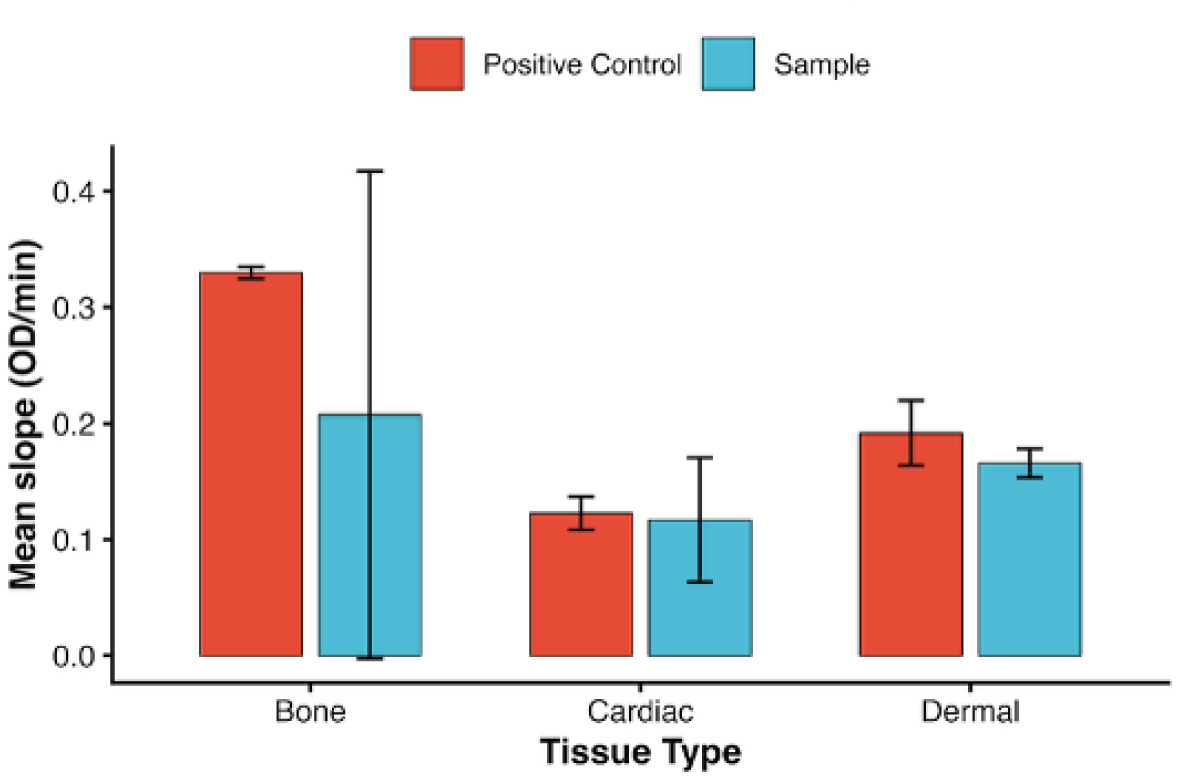
Pro-Collagen I Alpha 1 production across human cell lineages cultured on placental dECM or without it. Collagen production was quantified using a kinetic ELISA and compared between cells cultured on placental dECM and positive controls. Bars represent mean ± SD of biological replicates. Reproduced from Amurrio Zamora [22].

The greatest reduction in signal intensity between ECM and control conditions was observed for hFOB cells, whereas HCF cultures showed nearly identical values across conditions. Despite these trends, statistical analysis showed differences between ECM and positive control groups were not significant within the current experimental sample size.

Overall, these findings demonstrate that all three human cell lineages remained viable and continued to produce detectable levels of Pro-Collagen I Alpha 1 while cultured on dehydrated placental ECM scaffolds.

## Discussion

This study investigated whether a dehydrated placenta-derived ECM scaffold could support attachment and early adaptation across multiple human cell lineages while retaining biologically relevant cell-matrix interactions following processing. By using a combination of fluorescence imaging, quantitative morphometric analysis, and biochemical assessment, the present study demonstrates that placental ECM provides a permissive microenvironment capable of supporting the attachment and survival of dermal fibroblasts, cardiac fibroblasts, and osteoblasts. Additionally, the scaffold induced measurable, lineage-dependent changes in nuclear morphology, suggesting that processed placental ECM remains biologically active despite the tissue processing steps, including dehydration and terminal irradiation.

Fluorescence microscopy showed successful attachment and cytoskeletal organization across all three cell types cultured on the scaffold. The presence of nuclei and organized actin suggests that the ECM scaffold supports early cell survival and adhesion. This is consistent with the established role of ECM components in regulating structural integrity and cell signaling [25]. These findings also agree with previous studies that reported that amnion-derived biomaterials retain bioactive components capable of promoting cellular attachment and viability following tissue processing [15,26].

Biochemical analysis further supported the observation that all three cell types remained viable following culture on the placental ECM scaffold. Collagen is the most abundant structural protein in the ECM and is a major component of connective tissues [11]. Quantifying Pro-Collagen I Alpha 1 provided a functional measure of matrix-producing activity in cells cultured on the placental ECM scaffold. The production of Pro-Collagen I Alpha 1 was detected across all experimental groups. This indicated that cells maintained functional activity after attachment to the biomaterial. Collagen production was consistently lower under ECM conditions when compared to culture plastic controls, yet these differences did not reach statistical significance within the current sample size. These findings suggest that conventional biochemical assays alone may provide a limited view of early cell-material interactions, specifically when viability is preserved across experimental conditions.

Although all three cell types remained viable on the scaffold, quantitative morphometric analysis revealed significant differences in nuclear architecture between ECM and standard culture conditions. Overall, cells that were cultured on placental ECM exhibited increased nuclear eccentricity across all lineages, while circularity was significantly altered in HCF and hFOB cells. Nuclear morphology reflects the integration of extracellular mechanical and biochemical signals transmitted through the adhesions of the cell and ECM, the actin cytoskeleton, and the nuclear envelope. Together, these results indicate that the scaffold architecture actively influences cellular organization through mechanotransduction pathways rather than functioning as an inert support material as well as tissue-specific receptor interactions and growth factor signaling [27,28,29,30,31]. The dehydrated placental ECM therefore appears to modify the physical microenvironment experienced by adherent cells while preserving a substrate that supports cell attachment and viability. At the same time, the ability of the single processed ECM scaffold to support multiple unrelated human cell populations suggests that dehydration may reduce some aspects of tissue specificity while preserving a biologically supportive environment.

From this perspective, one of the main contributions of the present study is not the characterization of a placental ECM scaffold, but the demonstration that computational morphometric profiling can serve as a sensitive and quantitative approach for evaluating biomaterial performance. Conventional scaffold characterization often heavily relies on qualitative microscopy and endpoint viability assays, which may fail to detect subtle but biologically significant changes in cellular adaptation. The present workflow captures phenotypic responses that would likely remain undetected using standard approaches by combining automated deep learning segmentation with single-cell morphometric feature extraction [32,33,34].

Importantly, this analytical framework is not limited to placental ECM biomaterials. The image-analysis pipeline developed here could be applied to the evaluation of decellularized tissues, synthetic ECM and other regenerative biomaterials, providing an objective method for quantifying interactions between cells and biomaterials across diverse experimental systems. As regenerative medicine increasingly moves toward data-driven biomaterial design, integrating computational phenotyping with traditional biological assays may improve the ability to distinguish between scaffolds that merely support survival and those that actively influence cellular behavior. Several limitations should be considered when interpreting these findings. The relatively small number of biological replicates limited statistical power, particularly for the biochemical analysis. In addition, functional assessment was restricted to a single extracellular matrix marker, and future studies incorporating broader molecular and transcriptional endpoints would provide a more comprehensive characterization of scaffold-mediated responses. Variability associated with scaffold handling and the physical behavior of the dehydrated membrane within the culture wells may also have contributed to heterogeneity in cell distribution and downstream measurements. Future work should focus on expanding this computational framework by integrating additional morphometric descriptors, spatial analyses, and machine learning approaches capable of linking cellular phenotype to biomaterial properties. Application of these methods to alternative ECM processing strategies, chemically modified scaffolds, and three-dimensional culture systems may provide new insights into how scaffold architecture influences cell fate and tissue regeneration.

In conclusion, this study demonstrates that dehydrated placental ECM supports attachment and survival across multiple human cell lineages while inducing measurable changes in nuclear morphology. More broadly, these findings establish quantitative morphometric profiling as a sensitive and scalable tool for characterizing cell–biomaterial interactions, highlighting its potential to complement and extend traditional approaches used in regenerative medicine research.

## Author Contributions

Conceptualization, CAZ, AI, ND, and AT; methodology, CAZ, AI, FM, and AT; software, CAZ, and FM; formal analysis, CAZ and FM; investigation, CAZ; resources, AI, ND, AT, and FM; data curation, CAZ; writing—original draft preparation, CAZ; writing—review and editing, CAZ, AI, ND, AT, and FM; visualization, CAZ; supervision, AT and FM; project administration, AT, and FM. All authors have read and agreed to the published version of the manuscript.

## Funding

No external funding was supplied for this research.

## Institutional Review Board Statement

N/A

## Informed Consent Statement

N/A

## Data Availability Statement

The datasets presented in this article are not readily available because of industry collaboration and an ongoing study with membrane allografts. Requests to access the datasets should be directed to Catalina Amurrio Zamora at.

## Conflicts of Interest

A.I. and A.T. are paid employees of Axolotl Biologix.

